# The kidney protects against sepsis by producing systemic uromodulin

**DOI:** 10.1101/2021.01.08.425960

**Authors:** Kaice A. LaFavers, Chadi Hage, Varun Gaur, Radmila Micanovic, Takashi Hato, Shehnaz Khan, Seth Winfree, Simit Doshi, Ranjani N. Moorthi, Homer Twigg, Xue-Ru Wu, Pierre C. Dagher, Edward Srour, Tarek M. El-Achkar

**Affiliations:** Indiana University School of Medicine, Department of Medicine, Division of Nephrology and Hypertension, Indianapolis, IN; Indiana University School of Medicine, Department of Medicine, Division of Pulmonary Medicine, Indianapolis, IN; Southern Indiana Nephrology and Hypertension, Columbus, IN; Indiana University School of Medicine, Department of Anatomy, Cell Biology and Cellular Physiology, Indianapolis, IN; New York University, Departments of Urology and Pathology, and VA New York Harbor Healthcare System, New York, NY; Roudebush VA Medical Center, Indianapolis, Indiana; Indiana University School of Medicine, Department of Medicine, Division of Hematology and Oncology, Indianapolis, IN; Indiana University School of Medicine, Department of Microbiology and Immunology, Indianapolis, IN

## Abstract

Sepsis is a significant cause of mortality in hospitalized patients. Concomitant development of acute kidney injury (AKI) increases sepsis mortality through unclear mechanisms. While electrolyte disturbances and toxic metabolite buildup during AKI could be important, it is possible that the kidney produces a protective molecule lost during sepsis with AKI. We previously demonstrated that systemic Tamm-Horsfall Protein (THP, uromodulin), a kidney-derived protein with immunomodulatory properties, falls in AKI. Using a mouse sepsis model without severe kidney injury, we show that the kidney increases circulating THP by enhancing basolateral release of THP from medullary thick ascending limb cells. In sepsis patients, changes in circulating THP are positively associated with critical illness. THP is also found *de novo* in injured lungs. Genetic ablation of THP in mice leads to increased mortality and bacterial burden during sepsis. Consistent with the increased bacterial burden, the presence of THP *in vitro* and *in vivo* leads macrophages and monocytes to upregulate a transcriptional program promoting cell migration, phagocytosis and chemotaxis and treatment of macrophages with purified THP increases phagocytosis. Rescue of septic THP^-/-^ mice with exogenous systemic THP improves survival. Together, these findings suggest that through releasing THP, the kidney modulates the immune response in sepsis by enhancing mononuclear phagocyte function and systemic THP has therapeutic potential in sepsis.

**Significance Statement:** Sepsis is a significant contributor to kidney injury as well as morbidity and mortality worldwide. Specific therapies to improve outcomes in sepsis with kidney injury have largely been limited to symptom management and infectious agent control, in part because it is unclear how kidney injury increases sepsis mortality. This paper describes the identification of Tamm-Horsfall protein, previously known to protect in ischemic models of AKI, as protective in preclinical models of sepsis. It demonstrates how the loss of THP leads to decreased mononuclear phagocyte function and diversity, increased pathogen burden and decreased survival. THP also increases in sepsis without severe kidney injury and concentrates in injured organs. Further study of THP in sepsis could lead to novel sepsis therapeutics.

## Introduction

Sepsis is a leading cause of death and disability worldwide (1). During sepsis, an aberrant systemic host response to infection causes organ damage and failure (2). Acute kidney injury (AKI) occurs in approximately 20-30% of sepsis patients (3) where it doubles mortality (4, 5), though the underlying mechanisms driving this increase are unclear (6). While reduced filtration capability is likely contributing to decreased survival, the kidney may also produce and release a protective molecule(s) lost in AKI.

Tamm-Horsfall Protein (THP, Uromodulin) is produced in the kidney primarily by Thick Ascending Limb (TAL) cells (7, 8). The majority is secreted apically as a high molecular weight polymer (9), constituting the major protein component of urine (10). Urinary THP regulates the ion channel activity within the TAL and collecting duct (11–14), allowing it to modulate salt sensitivity and calcium/magnesium homeostasis (12, 15, 16), and also confers protection from kidney stones and urinary tract infections (17). A smaller proportion of THP is secreted, un-polymerized, into the interstitium and circulation (18) as well as the urine (19). Increased circulating THP is associated with protection against chronic kidney disease, cardiovascular events and all-cause mortality in patients independently of kidney function (6, 20, 21). Our studies in a knockout mouse (THP^-/-^) model revealed that THP protects against kidney injury (20, 22, 23) and systemic oxidative stress (24) and regulates the local and systemic innate immune response (18, 21). Based on these studies, we hypothesize that THP plays a protective role in sepsis. Since severe ischemic AKI causes acute systemic THP deficiency (24), increased mortality in sepsis with AKI severe enough to lead to THP deficiency could be partially due to the loss of a protective role of circulating THP.

We demonstrated that circulating THP increases in a mouse model of sepsis. In a sepsis patient cohort without severe AKI increasing circulating THP positively correlates with sustained critical illness. In a separate cohort with respiratory complications from sepsis, we found circulating THP can concentrate in the lungs. In a cecal ligation and puncture model of sepsis in THP^-/-^ mice, THP protects against sepsis mortality and treatment with a monomeric form of THP increases survival. THP acts on macrophages to enhance phagocytosis and bacterial clearance, likely a key mechanism underlying THP protection in sepsis. Our results support that circulating THP is a protective molecule released by the kidney during sepsis, and its deficiency worsens the prognosis.

## Materials and Methods

### Study approval

Plasma samples from sepsis patients were obtained after written informed consent was obtained, in accordance with the Declaration of Helsinki. Bronchoalveolar lavage samples from ARDS and control patients were obtained within the framework of the Indiana BioBank after written informed consent was obtained, in accordance with the Declaration of Helsinki. All mice were bred, housed, and handled in the Association for Assessment and Accreditation of Laboratory Animal Care–accredited animal facility of Indiana University School of Medicine. All animal procedures were performed in accordance with the protocol approved by the Institutional Animal Care and Use Committee at Indiana University and Veterans Affairs.

### Sepsis Cohort

Inclusion criteria for the cohort included proven or suspected infection in at least one site, organ dysfunction in one or more organ system as measured by the Sequential Organ Failure Assessment Score and two or more of the following: core temperature > 38 or < 36°C, heart rate greater than 90 beats per minute, a respiratory rate twenty ≥ breaths per minute or PaCO2 ≤ 32 mmHg or use of mechanical ventilation for an acute process, a white blood cell count ≥ 12,000/ml or ≤ 4,000/ml, or immature neutrophils greater than 10%. Patients were excluded if they were below eighteen years of age, if they were judged to be in a terminal state, if sepsis developed after a proven or suspected head injury that was clinically active at the time of admission, if more than 24 hours had passed since inclusion criteria were met, if they had stage 5 chronic or end stage kidney disease or if they were included in other experimental studies. After informed consent, all the patients enrolled into the study based on the criteria had approximately 100 µl of serum isolated at admission to ICU and 48 hours later. Additionally, the following was extracted from electronic medical records to calculate Sequential Organ Failure Assessment Scores: baseline demographics, comorbid diseases, presence of CKD and state, complete blood count, basic metabolic panel, liver function test, heart rate, systolic and diastolic blood pressures, body temperature, microbiology data and site of infection, Glasgow Coma scale rating, arterial blood gas values, supplementary oxygen and ventilator settings, urine outputs, urinalysis and need for renal replacement therapy.

### ARDS Cohort

Bronchoalveolar lavage (BAL) samples from ARDS patients with a clinical diagnosis of sepsis and control patients were obtained from a clinical BAL lab at Indiana University which has IRB approval to perform research on residual specimens (PI – Homer Twigg, M.D.). The lab receives clinical specimens from multiple physicians and hospitals for cellular differential analysis. Thus, the BAL technique was not standardized among subjects. ARDS samples for this study were selected based on the presence of respiratory failure, diffuse infiltrates on chest imaging, and a marked neutrophilic alveolitis on BAL analysis, and the diagnosis was adjudicated by chart review for these patients. Control subjects consisted of BAL from normal subjects participating in other studies in the PI’s lab.

### Mice

Animal experiments and protocols were approved by the Indianapolis VA Animal Care and Use Committee. Age matched 8-12 week-old Tamm-Horsfall protein (THP) knockout male mice (129/SvEv THP^-/-^) and wild type background strain (129/SvEv THP^+/+^) were used (25). Cecal ligation and puncture was performed according to accepted protocols (26). In brief, mice were anesthetized with isoflurane and placed on a homeothermic table to maintain core body temperature at 37°C. The cecum was exposed via a midline incision and ligated with sutures. The ligated cecum was perforated with a 25-gauge needle. A small amount of fecal material was expressed into the abdominal cavity prior to closure of the incision. All animals received a subcutaneous analgesic (buprenorphine, 0.03 mg/mL, administered at 0.1 mg/kg) at the end of surgery and at the beginning/end of each subsequent day prior to sacrifice. Animals treated with 0.5-0.7 mg/kg tTHP (purification described below) or vehicle (5% dextrose) were injected intravenously 1-2 hours after surgery. For bacterial burden analysis, organs were removed aseptically and homogenates prepared in cold sterile PBS using the PreCellLys tissue homogenizer. Serially diluted homogenates were plated on blood agar and incubated for 24 hours at 37° C before imaging and colony enumeration. Peritoneal fluid was harvested by washing the peritoneal cavity with sterile PBS before serial dilution and plating as described for tissue homogenates. Animals with evidence of cecal rupture were removed from analysis.

### Real time PCR

Mouse kidneys were Trizol extracted before Precellys homogenization (Bertin Instruments) per the manufacturer’s protocol, and the extracted RNA reverse transcribed as previously described (22). Real-time PCR was performed in an Applied Biosystems (AB) ViiA 7 system using TaqMan Gene Expression Assays. The following primers were used: THP: Mm00447649_m1, GAPDH: Mm99999915_g1. Cycling parameters were: 50°C for 2 min, 95°C for 10 min, 40 cycles of 95°C for 15 sec and then 60°C for 1 min. Expression values were normalized to GAPDH endogenous control and reported as fold change compared to control using the delta-delta CT method, per manufacturer’s instructions.

### Western Blot

Mouse kidneys were lysed in RIPA buffer before Precellys homogenization (Bertin Instruments) according to the manufacturer’s protocol. Samples were separated on a 4-12% Bis-Tris Bolt Gel (Invitrogen Life Technologies) and transferred to a 0.45 µm PVDF membrane. Blots were probed with a polyclonal sheep anti-mouse THP (LifeSpan Bio, LS-C126349) and imaged using Super Signal Femto West Maximum Sensitivity Substrate (ThermoFisher Scientific). Blots were stripped using Restore Western Blot Stripping Buffer (ThermoFisher Scientific) and probed with rabbit anti-mouse beta actin-HRP (Abcam, ab20272)

### Tamm-Horsfall protein purification

Tamm-Horsfall protein was purified from normal human urine according to the method of Tamm and Horsfall (10). THP was subsequently treated with 8M urea and resolved using size exclusion chromatography to recover a non-aggregated form in the range of 60-120 kiloDaltons (kDa), as described recently (18). The purified THP was kept in a 5% Dextrose water solution. In all *in vitro* experiments, the concentration of endotoxin was < 0.02 EU/ml, measured using Limulus Amebocyte Lysate (LAL) assay.

### Immuno-fluorescence confocal microscopy

Kidney sections (50 µm, fixed with 4% paraformaldehyde) were prepared for immuno- fluorescence confocal microscopy as described previously (22). Briefly, immunostaining was done at room temperature, overnight, with primary antibody (sheep anti-THP, R&D Systems AF5144) in PBS + 2% BSA + 0.1 % Triton X-100 and secondary antibody + FITC-phalloidin (Molecular Probes, D1306) + DAPI (Sigma, D9542) in the same buffer. After mounting on glass slides, stained sections were viewed with the Olympus Fluoview laser-scanning confocal microscope. Images were collected under X20 and X60 magnification. Quantitation of the cellular distribution of THP fluorescence in thick ascending limb tubules (identified by presence of THP staining and absence of phalloidin staining) was performed on high power fields (3-4 fields per section, 1-2 sections per mouse, 4 mice per experimental group) using ImageJ. Apical and basolateral membrane associated THP average intensity was measured along the appropriate border in a 3 pixel (1.17 µm) spanning region of interest using a brush tool. The ratio of basolateral membrane associated THP: apical membrane associated THP was calculated for each tubule.

### Preparation of Bone Marrow Derived Macrophages

Bone marrow derived macrophages were generated as described previously (27). Briefly, femurs and tibia from euthanized THP^+/+^ mice were harvested aseptically into sterile PBS/1% FBS. Excess muscle was removed, and the leg bones were severed proximal to each joint. A 25-gauge needle was used to flush each bone with cold sterile PBS/1% FBS. Cells were centrifuged for 10 min at 500 x g (room temperature). Supernatant was discarded and the cell pellet resuspended in RPMI 1640 with L-glutamine, 10% FBS and 1X pen/strep. Bone marrow cells were differentiated into macrophages by plating them at a density of 2-4 x 10^5^ cells/ml in RPMI 1640 with L-glutamine, 10% FBS and 1X pen/strep supplemented with 50 ng/ml M-CSF. Partial media changes were performed every three days after seeding. Macrophages were used for experiments within 7-10 days after seeding.

### Phagocytosis assay

BMDM were pre-treated with 1 µg/ml truncated hTHP (prepared as described above) or 5% dextrose for two hours before incubating with pHrodo Green E. coli BioParticles (Invitrogen, Cat #P35366l, diluted 1:500 from the manufacturer’s protocol) for 15 minutes. Cells were washed and harvested by treatment with Tryple for 10 minutes at 37° C and then centrifuged at 300 x g for 5 minutes at room temperature. Cells were resuspended in PBS/1% FBS and mixed with an equal volume of 1 µg/ml propidium iodide. Samples were kept on ice before being run on the Guava easyCyte flow cytometer. Control samples (unlabeled cells/no beads/no PI, cells/beads/no PI, cells/no beads/PI, beads alone) were used to determine live/dead gates and to determine the percentage of live cells which had taken up the fluorescent BioParticles.

### Single cell RNA-sequencing

BMDM were treated with 1 µg/ml truncated hTHP (prepared as described above) or vehicle for eighteen hours. Cells were harvested as above before being counted and resuspended in ice cold PBS/2% FBS/1 mM CaCl_2_.

For isolation of CD45^+^ cells from sham-operated kidneys of THP^+/+^ and THP^-/-^ mice, two mice were first subjected to sham surgery (abdomen exposed by midline incision and then sutured closed). Approximately 6-7 hours later, the animals were euthanized and kidneys removed and placed in Hank’s balanced salt solution on ice. Kidneys were homogenized with a Tissuemiser (Fisher Scientific) handheld homogenizer on ice by making 10 passes three times. The tissue slurry was incubated with collagenase type IA for 30 min at 37°C. The resulting suspension was passed through a 70 µm strainer and washed with PBS +1% fetal bovine serum (FBS) before pelleting at ∼1,600 x g for 10 min. To remove red blood cells and cellular debris the suspension was clarified on a Percoll gradient (28). At room temperature, the cell pellet was resuspended in 8 mL of 40% Percoll in PBS +1% FBS and layered on top of 3 mL of 70% Percoll in PBS +1% FBS at room temperature. The gradient was spun for 30 minutes at 900 x g at room temperature. The centrifuge brake was turned-off to not disturb the formed gradient and separated cells. Following the spin there was a fatty cellular debris layer at the top of the gradient, a fuzzy “buffy coat” at the interface between the 40% and 70% Percoll, a thin band of red blood cells just below that and a pellet of red blood cells. The buffy coat was removed and diluted to 15 mL with PBS +1% FBS and spun at 800 x g for 10 min at 4°C. After removing the supernatant, the pellet was resuspended in 1 mL PBS + 1% FBS. In preparation for labeling with CD for sorting by the flow cytometry facility, the suspension was centrifuged and resuspended in 100 µL to which CD16/32 antibody (BDBioscience) was added and incubated for 5 minutes at room temperature to block Fc gamma receptors. To label the cells with CD45 antibody, 10 µL (1.5 µg) APC conjugated CD45 antibody (clone 30F11, Miltenyi) was added to the 100 µL suspension and incubated for 30 minutes at 4°C. Cells were resuspended in PBS/1% FBS and mixed with an equal volume of 1 µg/ml propidium iodide. Cd45+ live cells were selected on the basis of PI and APC Cd45 staining. The sorted cells were collected on ice.

For both cell preparations, dead cells were removed using the Easy Sep Dead Cell (Annexin V) Removal Kit, according to the manufacturer’s direction. To remove calcium salts, the cells were washed twice with PBS/1% FBS and filtered (40 µm filter) before running on the Chromium 10X platform. Sequencing data was analyzed using the Seurat program in R Studio (29). Raw and processed data are available through the Gene Expression Omnibus repository under the following datasets: GSE171639 (sham-operated kidneys) and GSE189433 (bone marrow derived macrophages, to be accessed using the secure token “etmpmamifbqztkh” while the manuscript is in review).

### Measurement of human Tamm Horsfall Protein (THP)

Concentrations of human THP in human plasma, human bronchoalveloar lavage fluid and mouse serum of animals dosed with human THP were measured using an ELISA kit from Sigma Aldrich (Cat. #RAB0751) according to the manufacturer’s protocol. This ELISA has a detection range of 40.96 – 10,000 pg/ml with an intra-assay coefficient of variation of <10% and an inter-assay coefficient of variation of <12%. All samples that were compared directly were assayed at the same time to reduce variation due to a batch effect.

### Measurement of mouse Tamm Horsfall Protein (THP)

Concentrations of mouse THP in mouse serum were measured using an ELISA kit from LifeSpan Bio (Cat. # LS-F6973-1) according to the manufacturer’s protocol. This ELISA has a detection range of 0.156 – 10 ng/ml with an intra-assay coefficient of variation of <10% and an inter-assay coefficient of variation of <12%. All samples that were compared directly were assayed at the same time to reduce variation due to a batch effect.

### Measurement of human serum albumin (HSA)

Concentrations of human serum albumin in human plasma and bronchoalveloar lavage fluid were measured using an ELISA kit from Sigma Aldrich (Cat. #RAB0603) according to the manufacturer’s protocol. This ELISA has a detection range of 4.915 – 1200 ng/ml with an intra-assay coefficient of variation of <10% and an inter-assay coefficient of variation of <12%. All samples that were compared directly were assayed at the same time to reduce variation due to a batch effect.

### Measurement of Serum Creatinine

Concentrations of mouse serum creatinine were measured by isotope dilution liquid chromatography tandem mass spectrometry by the University of Alabama O’Brien Center Bioanalytical Core facility.

### Statistical Analysis

Values of each experimental group are reported as mean ± standard deviation unless otherwise indicated. All statistical tests were performed using GraphPad Prism software unless otherwise indicated. A two tailed t-test was used to examine the difference in means for continuous data. An F test was used to compare variances; in cases where the variances were significantly different (p < 0.05), a Welch’s correction was applied to the t-test. A paired t-test was used for samples from the same patient collected at different times. The Fisher’s exact test was used to determine differences between categorical variables. Simple linear regressions were used to determine relationships between two continuous variables. Statistical significance was determined at p< 0.05. Statistical outliers were defined using the Tukey’s fences methods. Kaplan-Meier survival curves were analyzed using the Gehan-Breslow-Wilcoxon test to determine statistical significance.

## Results

### Circulating THP increases in murine sepsis

We used the murine cecal ligation and puncture (CLP) model, which mimics sepsis pathophysiology and does not cause rapid-onset severe AKI in young mice (26). Circulating THP increased by 6 hours after CLP (5.99 ± 1.16 ng/ml to 9.54 ± 2.49 ng/ml, p = 0.0019, Figure 1A) and remained elevated in survivors (14.18 ± 8.35 ng/ml, p = 0.0113, Figure 1A). To identify the source of increased circulating THP, we measured mRNA transcript and protein in the kidney following CLP. When compared to sham-operated animals, CLP-operated animals did not show a significant difference in the *UMOD* transcript (1.06 ± 0.12 in sham versus 1.08 ± 0.16 relative transcript level in CLP animals, p = 0.8564, Figure 1B) or THP protein (1.05 ± 0.85 in sham versus 0.95 ± 0.35 relative protein level in CLP animals, normalized to levels of actin and expressed as fold change over mean, p = 0.8360, Figure 1C). We then investigated whether THP trafficking to the basolateral side of the Thick Ascending Limbs (TALs), which is expected to supply THP to the serum, is increased. In the inner stripe of the mouse kidney medulla, which has the highest density of TAL cells, we found an increase in the ratio of basolateral to apical THP (0.33 ± 0.11 vs. 0.26 ± 0.08 in CLP vs. sham, respectively p < 0.0001, Figure 1D). These results suggest that increased levels of circulating THP are due to a shift in trafficking of THP towards the basolateral side of the TAL within the medulla.

**Fig. 1.**
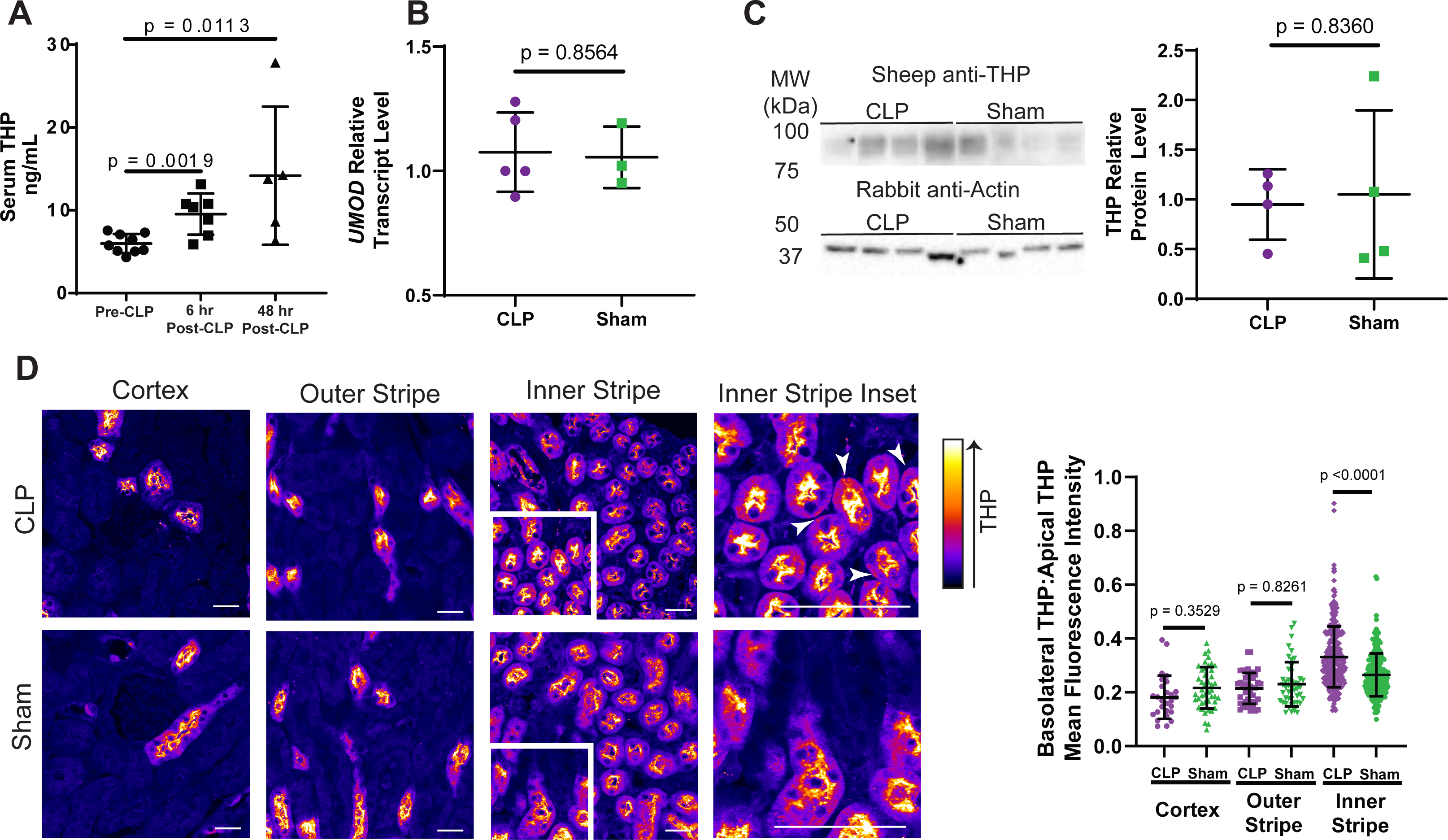
Circulating THP increases in sepsis due to increased basolateral trafficking within the inner stripe **(A)** Circulating THP levels increase following a mice model of sepsis (CLP) at both 6 and 48 hours after surgery **(B)** *UMOD* relative transcript levels are not significantly different in CLP versus sham operated mice 6 hours after surgery **(C)** THP relative protein levels (normalized to actin) are not significantly different in CLP versus sham operated mice 6 hours after surgery. Western Blot, left, quantitation, right. **(D)** Confocal microscopy of THP localization and intensity in CLP and Sham-operated kidneys 6 hours after surgery, left. Arrowheads indicate areas of high basolateral THP staining in the inner stripe of CLP-operated animals. The inner stripe of CLP-operated animals has increased basolateral THP:apical THP mean fluorescence intensity as compared to sham-operated animals. This phenomenon is restricted to the inner stripe, right. CLP – cecal ligation and puncture

### Circulating THP increases in clinical sepsis and concentrates in injured organs

We then enrolled a cohort of 26 patients from the Methodist Hospital Intensive Care Unit (Indianapolis) diagnosed with sepsis. Plasma and clinical parameters, used to calculate the Sequential Organ Failure Assessment (SOFA) score (30), were collected at admission and 48-hour follow-up. The SOFA score increases with organ dysfunction, damage and failure. Circulating THP was measured in plasma samples. The percent change in THP was positively correlated with follow-up (48 hour) SOFA score (R^2^ = 0.177, p = 0.0113, Figure 2A). Patients were stratified by follow-up SOFA score (Figure 2B). Those with SOFA score less than 6 had decreased plasma THP between admission and follow-up (49.3 ± 22.9 ng/ml versus 39.4 ± 19.0 ng/ml, p = 0.0200, Figure 2C). In patients with elevated (>6) follow-up scores, plasma THP remained high (45.7 ± 28.3 ng/ml versus 46.3 ± 26.5 ng/ml, p = 0.9453, Figure 2C). Proportions of patients with CKD in patients within each SOFA score group were similar (26.3% in the low SOFA group, 28.6% in the high SOFA group) and all had CKD stage 3, suggesting variations in serum THP levels are not due to impaired kidney function and ongoing critical illness is associated with higher circulating THP.

**Fig. 2.**
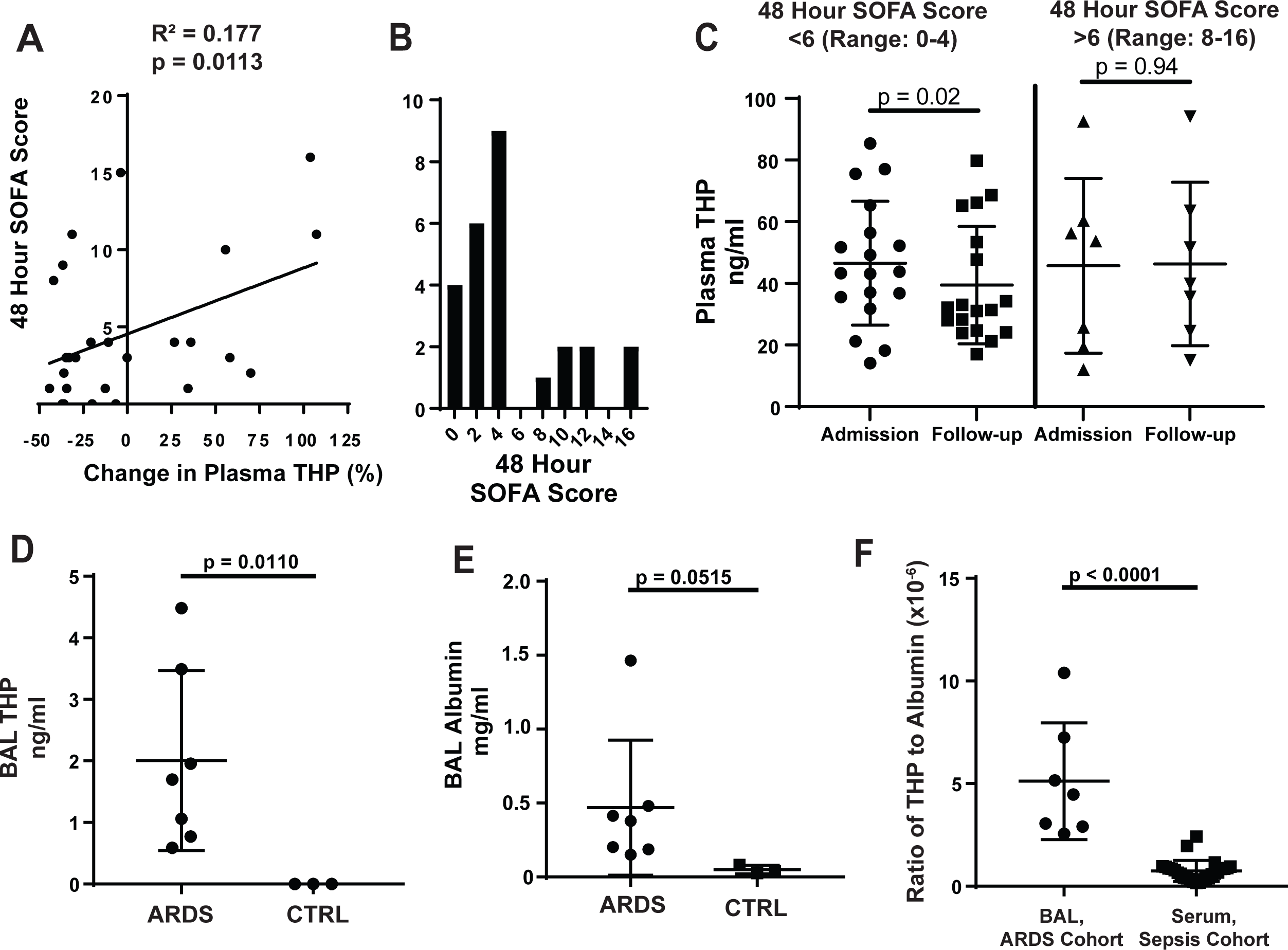
**Circulating THP increases with organ failure in sepsis and concentrates in injured organs** **(A)** In a cohort of patients admitted to the ICU with a diagnosis of sepsis, the percent increase in circulating THP from admission to 48-hour follow-up is positively correlated with 48-hour SOFA score (R^2^ = 0.177, p = 0.0323) **(B)** Histogram of 48 hour SOFA scores shows a natural break between patients with low organ dysfunction (0-4) or sustained organ dysfunction (8-16) **(C)** Plasma THP in patients with 48 hour SOFA score less than 6 drops significantly between admission and 48 hour follow-up, while patients with scores above 6 maintain levels of plasma THP **(D)** THP is found in the BAL fluid of patients with ARDS, but not in fluid from healthy controls (CTRL) **(E)** Albumin is elevated in patients with ARDS as compared to healthy controls **(F)** The ratio of THP:albumin is increased in the BAL fluid of the ARDS cohort as compared to the plasma of the sepsis cohort ICU – Intensive Care Unit, SOFA – Sequential Organ Failure Assessment, BAL – bronchoalveolar lavage, ARDS – acute respiratory distress syndrome

Since circulating THP has immunomodulatory and anti-inflammatory functions (17), we tested whether THP localized to distant injured organs in sepsis. Acute Respiratory Distress Syndrome (ARDS) is a rapidly progressing form of lung injury that can occur secondary to sepsis. Measurable THP was found in bronchoalveolar lavage (BAL) fluid of patients with ARDS (2.00 ± 1.46 ng/ml, Figure 2D, Table 2), but not healthy controls (p = 0.0110). In ARDS, disruption of alveolar barriers leads to interstitial edema with a protein-rich fluid (31), providing a potential mechanism for THP entry. To test this, we compared levels of human serum albumin (HSA) with THP. As shown previously (32), levels of HSA in the BAL fluid of ARDS patients increased compared to healthy controls (0.39 ± 0.40 mg/ml versus 0.05 ± 0.03 mg/ml, p = 0.0267, Figure 2E). THP and HSA levels in ARDS BAL fluid were positively correlated (Figure S1A, R^2^ = 0.6779, p = 0.0034), supporting the hypothesis that THP enters the lungs from the circulation following barrier dysfunction. In contrast, THP and HSA in plasma samples from the sepsis cohort were not significantly correlated (Figure S1B, R^2^ = 0.005668, p = 0.7147). To test whether THP is relatively concentrated in the lung compared to the circulation, we generated a ratio of THP to HSA concentrations for the BAL samples (ARDS cohort) and plasma samples (sepsis cohort). We found the ratio of THP:HSA is higher in the BAL samples (4.56 x 10^-6^ ± 2.92 x 10^-6^) compared to plasma samples (0.75 x 10^-6^ ± 0.51 x 10^-6^, p<0.0001), suggesting that THP could be concentrating in the lungs of ARDS patients (Figure 2F). These results demonstrate that the clinical response to sepsis without significant kidney injury is to increase levels of THP in the circulation and, potentially, damaged organs. Since previous work with THP suggested that it would have a protective role in the setting of infection, we hypothesized that higher levels of circulating THP are a protective mechanism rather than a causative agent of organ damage and dysfunction and tested this hypothesis in THP^-/-^ mice.

**Table 1.** Sepsis cohort characteristics at admission and follow-up

| Patient Characteristics (n = 26) |  |  |  |
| --- | --- | --- | --- |
| **Sex** |  |  |  |
| Male | 42% |  |  |
| Female | 58% |  |  |
| **Age (years)** | 57.4 ± 15.1 |  |  |
| **Race** |  |  |  |
| Caucasian | 88% |  |  |
| African | 12% |  |  |
| **Comorbidities** |  |  |  |
| Diabetes | 42% |  |  |
| COPD | 35% |  |  |
| CKD | 27% |  |  |
| HTN | 73% |  |  |
| Liver Disease | 11% |  |  |
| CHF | 27% |  |  |
| Malignancy | 15% |  |  |
| **Baseline Serum Creatinine (mg/dL)** | 0.82 ± 0.28 |  |  |
| **Baseline eGFR (mL/min)** | 100 ± 41.8 |  |  |
| **Infection source** |  |  |  |
| Lung | 12 (46%) |  |  |
| Abdomen | 6 (23%) |  |  |
| Heart | 2 (7.8%) |  |  |
| Skin | 2 (7.8%) |  |  |
| Blood | 3 (11.5%) |  |  |
| Urinary Tract | 1 (3.8%) |  |  |
| **Microbe identity** |  |  |  |
| Gram negative bacteria | 10 (38.4%) |  |  |
| Gram positive bacteria | 8 (30.8%) |  |  |
| Fungus | 1 (3.8%) |  |  |
| Unknown | 7 (27.0%) |  |  |
| **Required RRT** | 3 (11.5%) |  |  |
|  | **Admission** | **Follow-up** | **p-value** |
| **Plasma THP (ng/mL)** | 48.3 ± 23.9 | 45.8 ± 30.5 | 0.5108 |
| **SOFA Score** | 4.69 ± 2.78 | 4.58 ± 4.67 | 0.2092 |
| **Serum Creatinine (mg/dL)** | 1.48 ± 1.27 | 1.14 ± 0.873 | 0.2887 |
| **eGFR** | 80.1 ± 68.4 | 96.1 ± 73.9 | 0.0199 |

**Table 2.** ARDS cohort characteristics at admission and follow-up

| **Patient Characteristics (n = 7)** |  |
| --- | --- |
| **Sex** |  |
| Male | 14% |
| Female | 86% |
| **Age (years)** | 57.1 ± 7.88 |
| **Race** |  |
| Caucasian | 86% |
| African | 14% |
| **Comorbidities** |  |
| Diabetes | 14% |
| CKD | 14% |
| HTN | 71% |
| **Admission Serum Creatinine (mg/dl)** | 1.11 ± 0.598 |
| **Serum Creatinine on day of BAL (mg/dl)** | 0.734 ± 0.322 |

### THP enhances phagocytosis and reduces bacterial burden in sepsis

To gain further insight about the role of THP in sepsis, we performed CLP surgery in THP^-/-^ and THP^+/+^ mice (25) and found THP^-/-^ mice had significantly increased mortality (median survival of 1 versus 2 days, p = 0.0148, Figure 3A), establishing that THP is protective in sepsis. We propose that increased circulating THP is an important response from the kidney to control systemic infection. We compared serum creatinine levels 6 hours after CLP surgery for THP^+/+^ and THP^-/-^ mice and found that levels were not increased in either group (0.77 ± 0.15 µg/ml and 1.05 ± 0.13 µg/ml, respectively) compared to sham-operated animals of the same genotype (0.67± 0.09 µg/ml and 0.85 ± 0.13 µg/ml, respectively, p = 0.6908 for THP^+/+^, p = 0.1733 for THP^-/-^, Figure 3B), suggesting that early mortality seen in the CLP animals is not likely due to a rapidly progressing kidney injury. Given THP’s role in modulating innate immune function, we hypothesized that defects in the innate immune response could be responsible for the early mortality seen in THP^-/-^ mice.

**Fig. 3.**
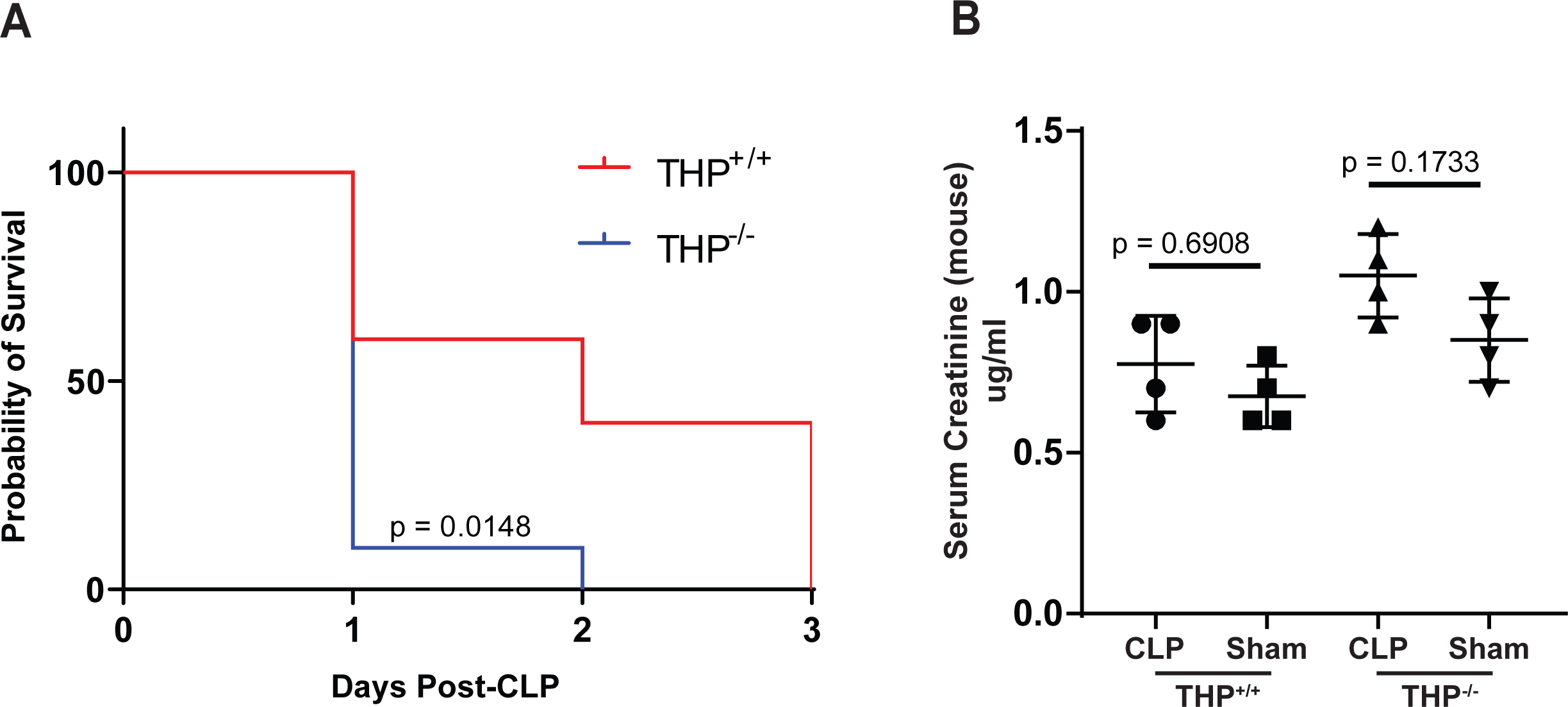
THP protects in a mouse model of sepsis **(A)** THP^-/-^ mice have increased mortality following CLP as compared to THP^+/+^ mice **(B)** Neither THP^+/+^ or THP^-/-^ mice have significantly elevated serum creatinine 6 hours post-surgery as compared to sham-operated animals

We previously showed THP^-/-^ mice have decreased number and function of kidney mononuclear phagocytes (MNP, (18)), leading us to hypothesize that THP enhances the function of MNP to protect against sepsis. To gain unbiased insights on the role of THP in MNP activation, we treated mouse bone marrow-derived macrophages (BMDM) from THP^+/+^ mice with 1 µg/ml of a truncated, non-polymerizing form of THP (tTHP), isolated from human urine which more accurately represents the monomeric form of THP found in the kidney interstitium and circulation (18) or vehicle before preparing cells for single cell RNA sequencing. BMDM were found to be a heterogenous population which can be sorted into 8 clusters (Figure 4A, left). Cluster 2 expands with tTHP treatment with a log-adjusted fold change of approximately 1.5 (Figure 4A, right). To predict the phenotype associated with macrophages within Cluster 2, we identified the top 15 markers for each cluster (heatmap, Supplemental Figure 2). Within Cluster 2, many of the transcripts fell into the functional categories of cell migration and phagocytosis (Figure 4B, (33–38)). Finally, we did a pathway analysis and found that treatment with tTHP increased expression of genes involved in ribosome function (biogenesis, ribonucleoprotein complex biogenesis, rRNA processing, rRNA metabolic processes, ribosomal small subunit biogenesis) as well as response to external stimuli, ATP metabolic processes, apoptosis and leukocyte/cell chemotaxis (Figure 4C). Together, these results suggest that treatment of BMDM with tTHP increases their metabolic activity and ability to respond or chemotax in response to external stimuli.

**Fig. 4.**
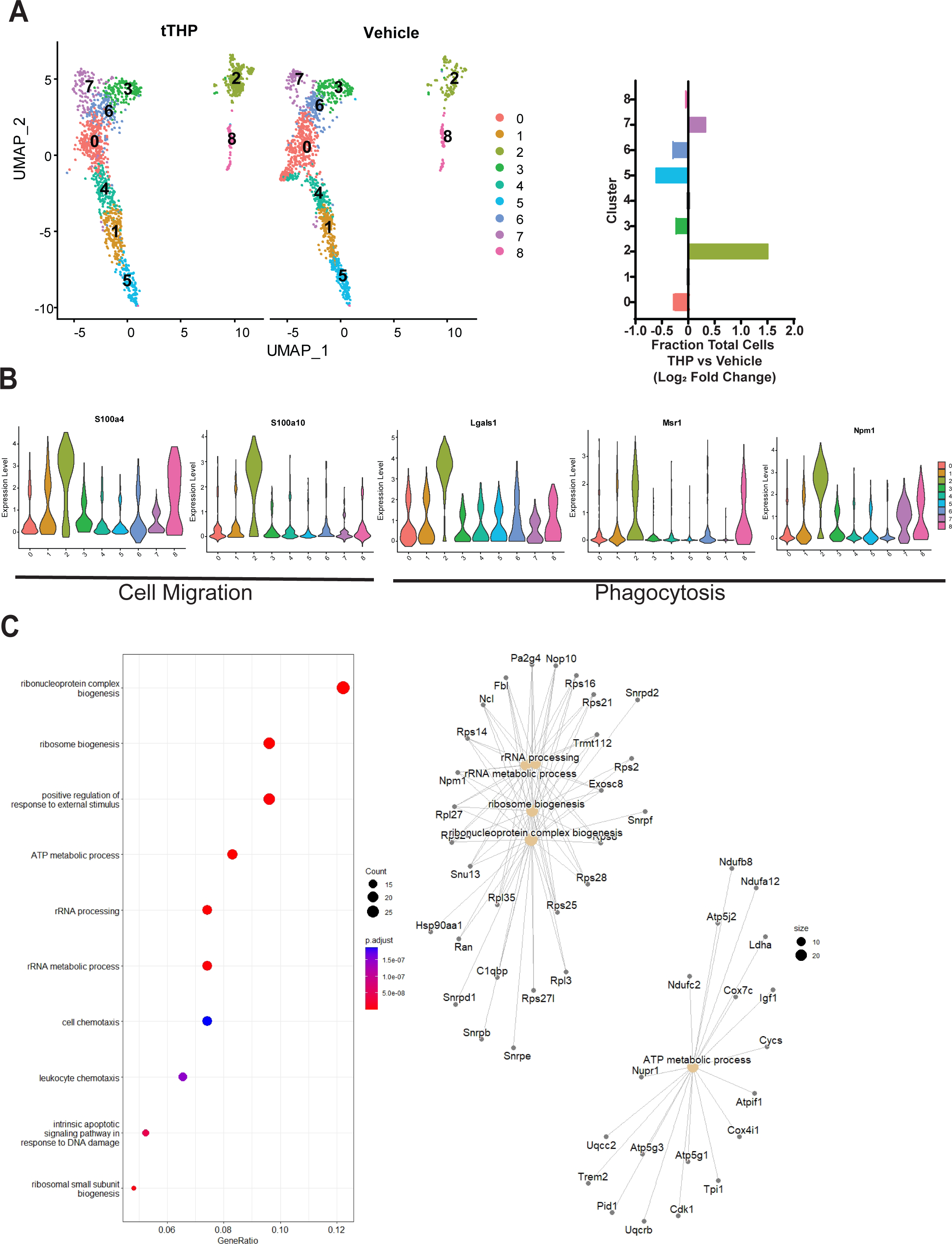
Treatment of Bone Marrow Derived Macrophages with tTHP leads to expansion of a subset of macrophages primed for phagocytosis **(A)** Single cell RNA sequencing of tTHP and vehicle-treated mouse BMDM reveals eight distinct clusters, left. The relative proportion of cluster two increases with THP treatment, right. **(B)** Markers of cluster 2 have roles in cell migration, left, and phagocytosis, right. Violin plots for representative transcripts show relative expression of markers in cluster 2 compared to other clusters. **(C)** Reactome pathway analysis of single cell RNA-sequencing data shows upregulation of specific pathways in tTHP-treated bone marrow derived macrophages (left) and the networks of genes contributing to these pathways (right) tTHP = truncated human THP

We performed a similar single cell RNA-sequencing experiment on Cd45^+^ cells isolated from kidneys of sham-operated THP^+/+^ and THP^-/-^ mice to perform a baseline characterization of MNP *in vivo*. After clustering and labeling the cells according to the ImmGen database((39), Figure 5A), we subset the MNP. Further identification of these cells showed that THP^-/-^ mice have decreased macrophage diversity and are lacking the large numbers of Ly6C^hi^ monocytes found in THP^+/+^ animals (Supplemental Figure 3). Ly6C^hi^ monocytes are known for their anti-microbial function and demonstrate high capability for phagocytosis (40). Many of the genes upregulated in response to tTHP treatment of BMDM involved in cell migration and phagocytosis were also increased in THP^+/+^ mice (Figure 5B). The pathway analysis of this scRNA-seq data showed increased expression of genes involved in lipid localization and phosphorylation that were also connected with immune cell migration and chemotaxis. Together with our previous findings that THP^-/-^ mice have impaired phagocytosis within the kidney (18), this suggested that THP increases the presence of MNP primed for chemotaxis and phagocytosis.

**Fig. 5.**
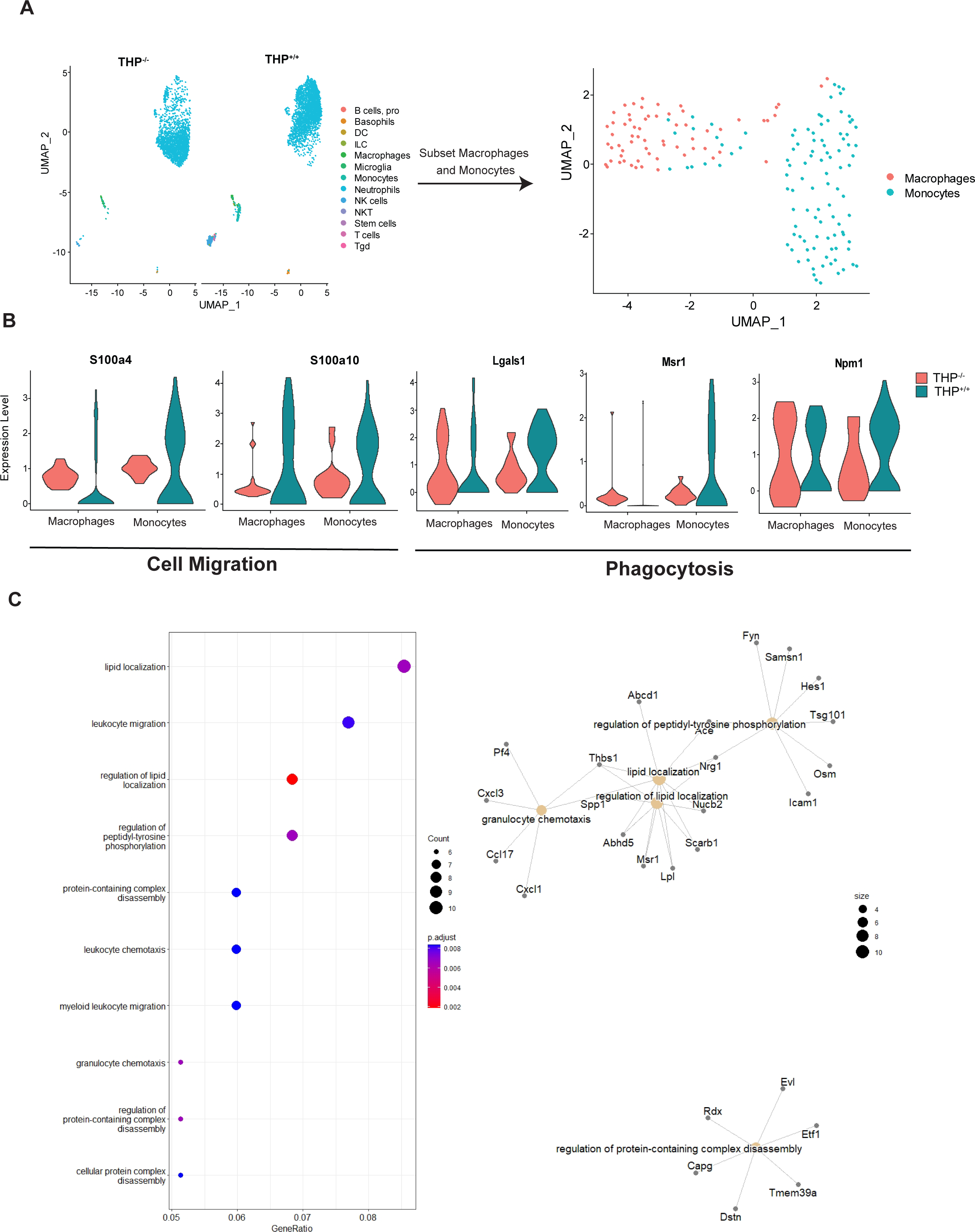
Macrophages and monocytes isolated from THP^+/+^ kidneys have activated transcriptional programs for cell migration and chemotaxis as compared to those from THP^-/-^ kidneys **(A)** Single cell RNA sequencing of Cd45^+^ cells isolated from sham-operated THP^+/+^ and THP^-/-^ kidneys reveals a heterogenous population of immune cells. From this, a population identified using the ImmGen database as macrophages and monocytes can be subset (right) **(B)** Violin plots of selected transcripts identified in the scRNA-seq of tTHP-treated BMDM macrophages which are associated with cell migration and phagocytosis are decreased in the macrophages and monocytes of THP^-/-^ mice compared to THP^+/+^ **(C)** Reactome pathway analysis of single cell RNA-sequencing data shows upregulation of specific pathways (left) in THP^+/+^ mice and the networks of genes contributing to these pathways (right)

Since MNP are important effectors of the innate immune response during infections, we hypothesized that THP enhances pathogen clearance at the organ level in sepsis by stimulating MNP activity. Since CLP induces a peritoneal infection, we examined bacterial burden in peritoneal fluid and abdominal organs six hours following surgery (Figure 6A). THP^-/-^ mice have an increased number of colony forming units (CFU) of bacteria within the spleen (1.87 ± 5.01 versus 17.6 ± 18.6 CFU/mg tissue, p = 0.0136), liver (0.668 ± 0.805 versus 1.74 ± 0.697 CFU/mg tissue, p = 0.0352) and peritoneal fluid (3667 ± 4381 versus 25,125 ± 14,631 CFU/ml, p = 0.0003, Figure 6A). These results support the key role of THP in bacterial clearance through enhancing MNP activity. This defect in clearance, manifesting early during infection, could explain the premature mortality of THP^-/-^ mice, and other states of THP deficiency, such as AKI (24), since the timing of antibiotic intervention and subsequent decrease in bacterial levels is a predictor of mortality in bacterial sepsis (41).

To test whether THP could be mediating this effect on MNP function directly, we treated BMDM with tTHP and measured uptake of fluorescently labeled BioParticles using flow cytometry. tTHP-treated macrophages have significantly increased phagocytosis compared to vehicle-treated macrophages (1 ± 0.279 versus 1.446 ± 0.36-fold change over vehicle, p = 0.0151, Figure 6B), supporting a direct enhancement of phagocytic activity by THP.

**Fig. 6.**
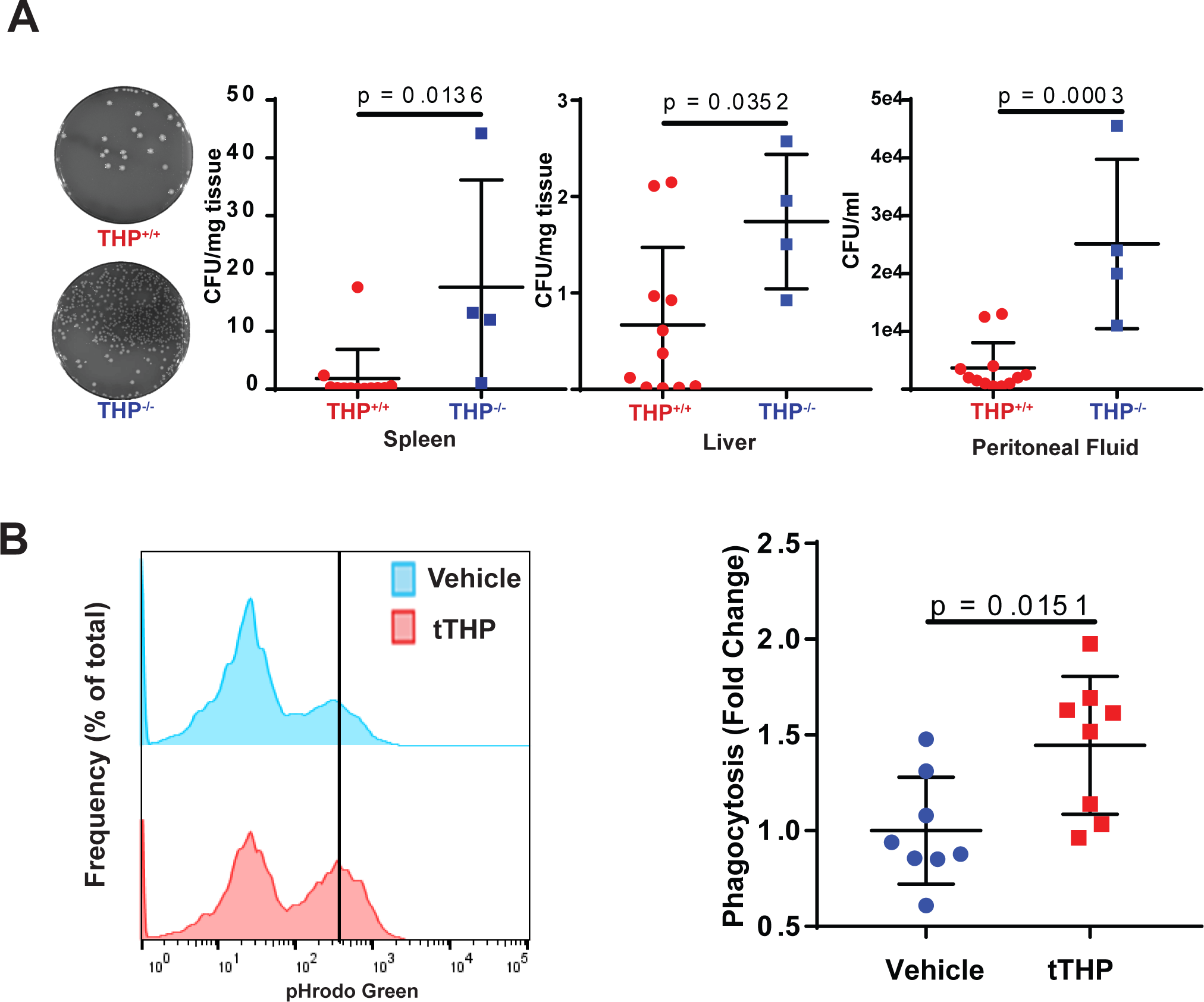
THP decreases bacterial burden *in vivo* sepsis and enhances phagocytosis *in vitro* **(A)** THP^-/-^ mice have an increased bacterial burden in the spleen, liver and peritoneal fluid (right) as compared to THP^+/+^ mice 6 hours following CLP surgery. Representative blood agar plates shown (left). **(B)** Treatment of BMDM with tTHP increases phagocytosis of pHrodo green fluorescently labeled BioParticles. Representative histogram of vehicle and tTHP treated macrophages, left, solid line indicates gates used for positive cells tTHP = truncated human THP

### THP treatment restores normal sepsis mortality in THP^-/-^ mice

We previously showed tTHP administered systemically in mice reaches kidney macrophages (18). We hypothesized that treatment of THP^-/-^ mice with tTHP may restore the protective systemic effects of THP and improve survival following sepsis.

THP^+/+^ and THP^-/-^ mice were injected intravenously with tTHP (THP^-/-^ only, dosed at approximately 0.5 mg/kg based on previous efficacy studies) or vehicle (both) after CLP surgery. Treatment of THP^-/-^ mice with tTHP restored survival to the levels of vehicle-treated THP^+/+^ mice, while vehicle-treated THP^-/-^ mice showed a significant increase in mortality (Figure 7).

**Fig. 7.**
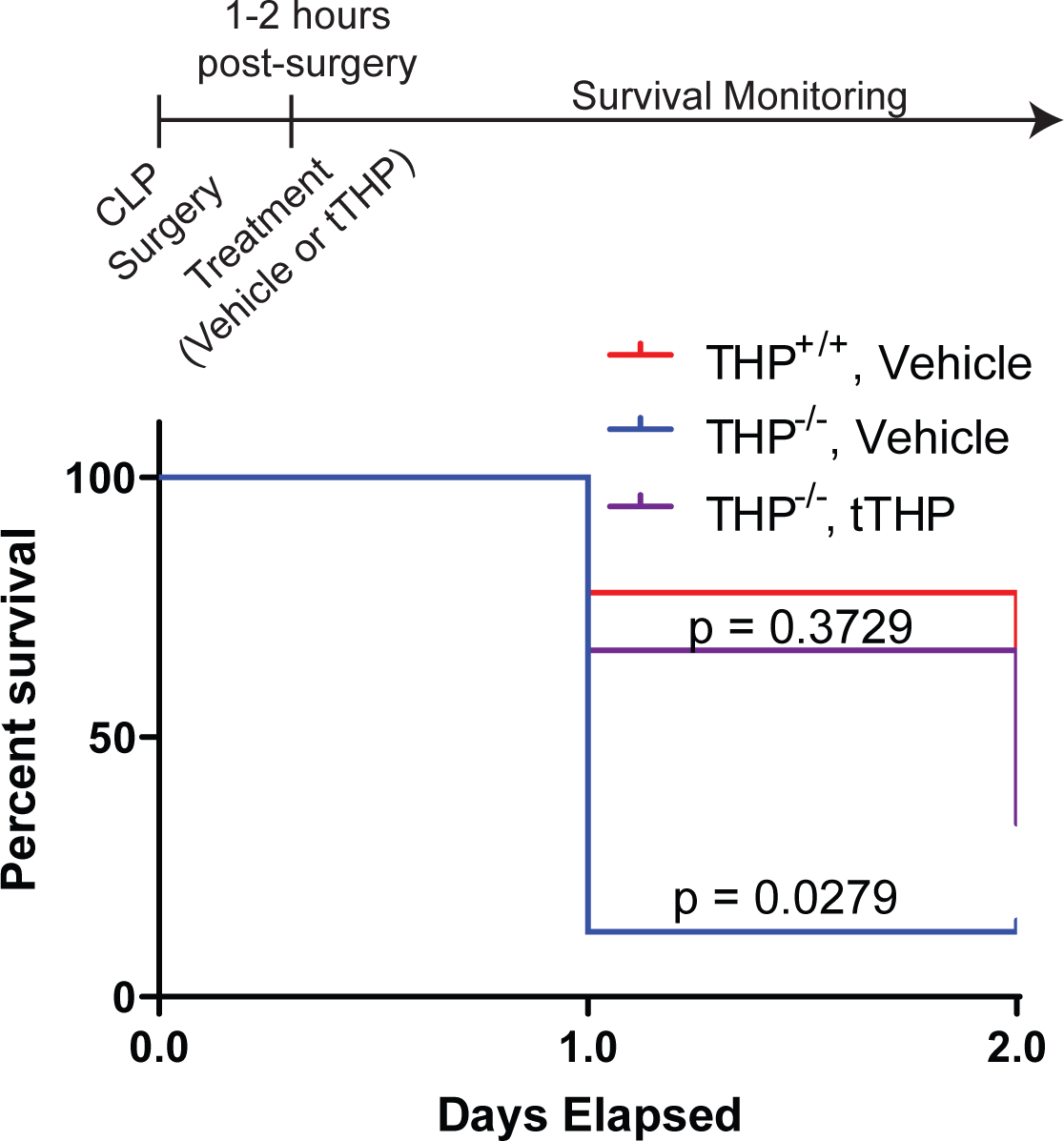
Treatment of THP^-/-^ mice with tTHP after CLP surgery restores survival to levels of THP^+/+^ mice tTHP = truncated human THP

## Discussion

Circulating THP increases in animal models of sepsis, as well as patients with sepsis and persistent critical illness. In mice, this is due to enhanced basolateral trafficking of THP protein within the medulla. Circulating THP also concentrates in damaged lungs of ARDS patients. The absence of THP worsens prognosis since THP^-/-^ mice had increased mortality compared to THP^+/+^ counterparts. Treatment of macrophages with THP expands cells characterized by expression of transcripts required for cell migration and phagocytosis and increases phagocytosis *in vitro*. Consistent with these findings, THP^-/-^ mice exhibit increased bacterial burden after CLP. Rescue of THP deficiency with systemic exogenous THP improves survival. Our findings support that THP protects in sepsis by increasing the phagocytic function of mononuclear phagocytes and reducing the microbial burden in the course of the disease.

The increased basolateral trafficking of THP within the inner stripe provides both a plausible mechanism for increased circulating THP in sepsis and an intriguing parallel to other injury settings. Basolaterally released THP increases during the recovery phase of AKI caused by ischemia repurfusion injury (22) and is known to have pleiotropic effects (17) on both local and distant targets. It is of particular interest that the increase in basolateral trafficking occurs mainly within the inner stripe of the medulla which contains the highest concentration of TALs. We have proposed previously that TALs act as a sensor to detect and respond to conditions which lead to damage of more vulnerable S3 segments, located predominantly in the outer stripe (42), and have demonstrated that THP mediates a protective crosstalk between TALs and S3 segments in the setting of AKI (20).We expect that basolateral trafficking of THP in the setting of sepsis is a protective measure by the kidney in response to systemic illness, consistent with increased circulating THP observed in both the CLP animal model and human sepsis patients. When THP production is increased in conditions of systemic illness and inflammation, the kidney is contributing to the activation and polarization of mononuclear phagocytes required to fight infection.

The dynamic changes in THP levels have important implications for the usefulness of measuring circulating THP. In healthy patients without kidney disease, serum THP is not associated with kidney function (43, 44), and there is evidence that serum and urine uromodulin are independently regulated (19, 45). Although it was proposed that serum THP reflects nephron mass and should remain unchanged without kidney injury (46), our data support that circulating THP is dynamically regulated in response to systemic stress such as infections. Therefore, we propose that future studies should include serial measurements of circulating THP, which may be quite informative in the setting of acute illness.

Sepsis is caused by factors related to the particular invading pathogen(s) and to the immune system of the host. By serving an immunomodulatory role in enhancing phagocytosis of macrophages, circulating THP controls the pathogen factors (i.e. their burden), thereby ameliorating the prognosis of recuperation and survival. However, we expect that THP’s beneficial effects in sepsis extend beyond engulfment of pathogens. Improving phagocytic function will likely also increase clean-up of damaged/dying cells and neutrophil-derived NETs (47), further improving survival outcomes. This vigilant cleanup of cell debris by active phagocytes is essential to recovery, as apoptotic cells contribute to organ dysfunction or failure. Other previously described functions of THP might also improve survival in sepsis, including inhibiting release of cytokines/chemokines that could promote a maladaptive immune response (12, 14, 17, 37) or acting as a cytokine trap (37). We have also recently demonstrated that THP reduces renal and systemic oxidative stress via inhibition of TRPM2 channel (17). This is of importance since oxidant injury is an important driver of sepsis pathology (39).

Our findings have particular significance in sepsis complicated by AKI. We recently demonstrated that AKI induces systemic THP deficiency (24). In sepsis with AKI, the loss of systemic THP could explain the increased mortality observed and provide a rationale for future studies to investigate therapeutic strategies to replete systemic THP. The successful rescue of THP^-/-^ mice with exogenous THP paves the way for developing these studies. Future studies on the pharmacokinetics and pharmacodynamics of THP replacement are needed, which could elucidate whether pharmacological augmentation of systemic THP could benefit sepsis without AKI or THP deficiency.

Our studies on the effect of THP on MNP signaling and function *in vivo* and *in vitro* present novel insights on THP’s immunomodulatory function. The decrease in macrophage density and Ly6C^+^ monocytes is consistent with our previous studies showing a relative deficiency in the CD11b^hi^, F4/80^lo^ population. Our study suggests that THP may have a role in MNP migration and phagocytosis. Our findings *in vitro* on BMDM were strikingly concordant with the *in vivo* studies and identified a transcriptomic signature that promotes macrophage activation, phagocytosis and chemotaxis. Future studies are needed to identify a particular receptor of THP on macrophages. Previous studies suggested THP interacts with the inhibitory neutrophil receptor sialic acid-binding Ig-like lectin-9 (Siglec-9), which could play a role in regulating phagocytosis (48). Ongoing studies are underway to determine if such interaction also occurs on MNPs or whether activation occurs through a different molecule.

Our study has limitations. The animal studies were performed in a constitutive knockout model where the mice could have adapted to the loss of THP. Salt wasting previously described in THP^-/-^ mice due to inactivation of the NKCC2 channel in TAL cells (13) could have contributed to the worse outcomes observed in these mice with sepsis. However, the rescue and improved survival with systemic THP argues against a major contribution of such an effect. The sepsis cohort has a small number of patients and the findings need to be reproduced in a larger study. The measurements of THP on admission to the ICU do not include baseline levels or data addressing genetic variability in the *UMOD* promoter, an important driver of THP production by the kidney (49). In this context, the direction of change is more relevant than absolute levels. Since the bronchoalveolar lavage samples were collected from a biobank, we did not have serum samples or lavage volumes which would enable us to directly compare the level of THP in the blood and BAL fluid from the same patients. The mechanisms of how THP is concentrated in the lungs are not clear and require further investigation. Our results could be the basis of future studies of THP levels in BAL fluid as a biomarker for sepsis and lung injury. Since isolating THP from mouse urine is not feasible, all treatments were done with THP isolated from human urine. As such, it is difficult to compare concentrations used with physiological levels as it is both possible and likely that the human form of the protein interacts with mouse receptors and cells with different binding kinetics as it does the mouse form.

In conclusion, we demonstrated that THP positively affects survival of septic animals, primarily through its immunomodulatory role on mononuclear phagocyte activity. These studies highlight the protective role of the kidney in sepsis, which is explained, in part, by its ability to increase the systemic release of THP. We believe that this work could pave the foundation for therapeutic use of THP in sepsis, a condition that remains an unmet medical need.

## Author Contributions

KL, RM, ES and TEA designed, analyzed and interpreted experiments and wrote the manuscript. TH and PD analyzed data and edited the manuscript. SD and RNM analyzed data. Transcriptomic data acquisition and analysis was performed by KL and SW. Protein biochemistry experiments were performed by KL and RM. Experiments in the tissue culture models were performed by KL. Animal studies were conducted by KL, SK, TEA, RM and TH. Collection of human cohort specimens and data was performed by CH, VG and HT.

Supplemental Figure 1

**(A)** Distribution of SOFA scores at 48 hours in the sepsis cohort

**(B)** Levels of albumin in BAL fluid from ARDS patients show a trend toward increased levels compared to CTRL patients.

**(C)** Levels of circulating THP are not significantly correlated with levels of circulating albumin (R^2^ = 0.005668, p = 0.7147) in the sepsis cohort BAL – bronchoalveolar lavage, ARDS – acute respiratory distress syndrome

Supplemental Figure 2

**(A)** Heat map of the top three markers defining each cluster of bone marrow derived macrophages

**(B)** Levels of human THP in the serum of THP^-/-^ mice treated with human tTHP

Supplemental Figure 3

Predicted cellular identities (ImmGen) of macrophages and monocytes isolated from

THP^-/-^ and THP^+/+^ kidneys.

## Acknowledgements

We acknowledge the Indiana Biobank for assistance obtaining the bronchoalveolar lavage samples (https://indianabiobank.org) and the University of Alabama O’Brien Core for serum creatinine measurements.

## Notes

**Funding:** This work was supported by The National Institute of Diabetes Digestive and Kidney Disease (NIDDK: 1R01DK111651 and P30DK079312 to TME), a Veterans Affairs Merit Award to TEA, and an American Society of Nephrology Ben J. Lipps award, T32 fellowship (5T32DK120524-02) and K99 award (1K99DK127216-01A1) to KL.

### Competing Interest Statement

TEA and RM have applied for a patent related to this work titled Modified Tamm-Horsfall Protein and related compositions and methods of use, publication number US 2018/0305420 A1.

### Summary of Updates

Additional analysis of scRNA-seq data in figures 4 and 5. Change in order of figures. Changes were made to the text to reflect the new data and order.

