## Supplementary figures and images for "The kidney protects against sepsis by producing systemic uromodulin"

### Supplemental Figure 1

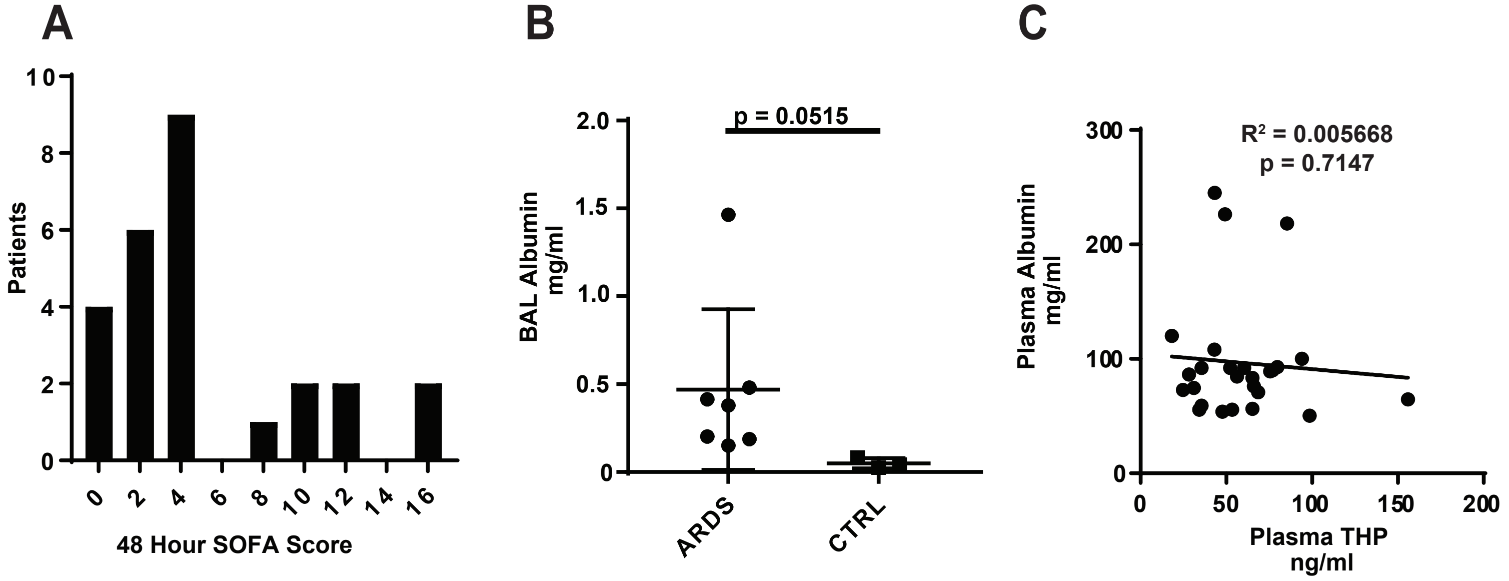

### Supplemental Figure 2

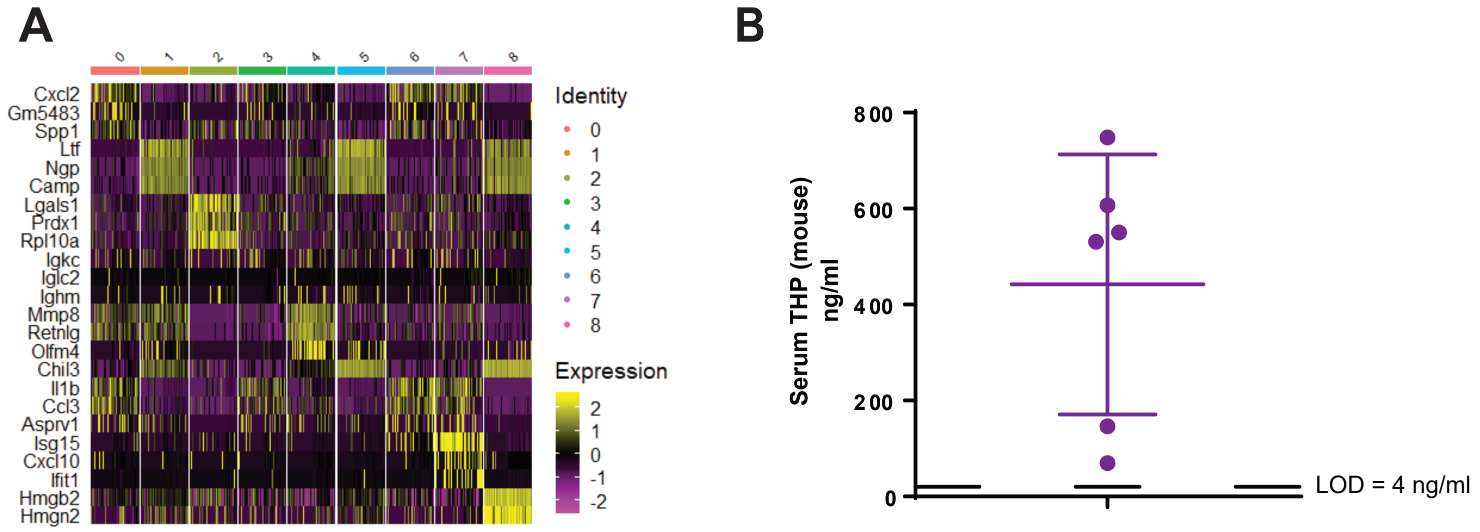

### Supplemental Figure 3

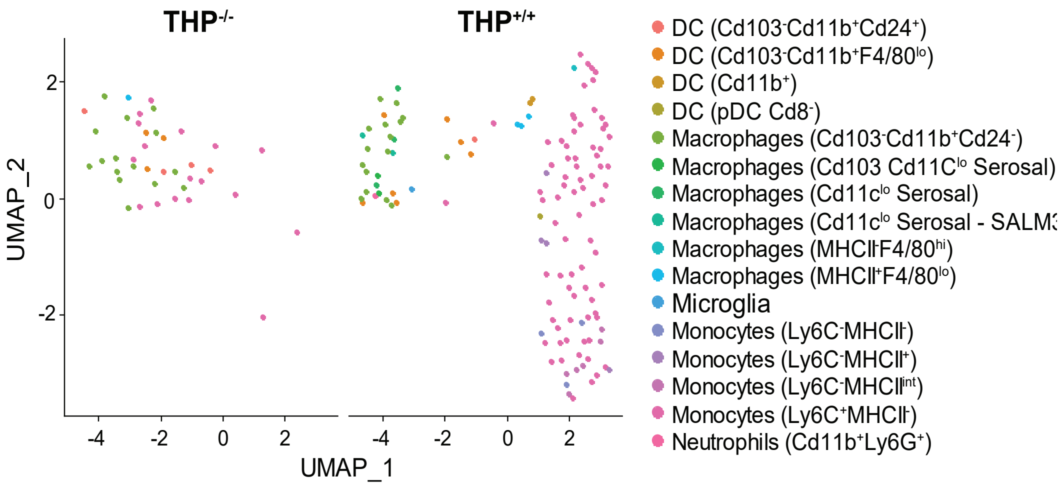
